# Visual processing speed is linked to functional connectivity between right frontoparietal and visual networks

**DOI:** 10.1101/2020.11.11.378406

**Authors:** Svenja Küchenhoff, Christian Sorg, Sebastian Schneider, Oliver Kohl, Hermann J. Müller, Kathrin Finke, Adriana L. Ruiz-Rizzo

## Abstract

Visual information processing requires an efficient visual attention system. The neural theory of visual attention (TVA) proposes that visual processing speed depends on the coordinated activity between frontoparietal and occipital brain areas. Previous research has shown that the coordinated activity between (i.e., functional connectivity, ‘inter-FC’) cingulo-opercular (COn) and right-frontoparietal (RFPn) networks is linked to visual processing speed. However, evidence for how inter-FC of COn and RFPn with *visual* networks links to visual processing speed is still missing. Forty-eight healthy human adult participants (27 females) underwent resting-state (rs-)fMRI and performed a whole-report psychophysical task. To obtain inter-FC, we analyzed the entire frequency range available in our rs-fMRI data (i.e., 0.01-0.4 Hz) to avoid discarding neural information. Following previous approaches, we analyzed the data across frequency bins (Hz): Slow-5 (0.01-0.027), Slow-4 (0.027-0.073), Slow-3 (0.073-0.198), and Slow-2 (0.198-0.4). We used the mathematical TVA framework to estimate an individual, latent-level visual processing speed parameter. We found that visual processing speed was negatively associated with inter-FC between RFPn and visual networks in Slow-5 and Slow-2, with no corresponding significant association for inter-FC between COn and visual networks. These results provide first empirical evidence that links inter-FC between RFPn and visual networks with the visual processing speed parameter. These findings suggest a direct connectivity between occipital and right frontoparietal, but not frontoinsular, regions, to support visual processing speed.

**Significance statement:** An efficient visual processing is at the core of visual cognition. Here, we provide evidence for a brain correlate of how fast individuals process visual stimuli. We used mathematical modeling of performance in a visual report task to estimate visual processing speed. A frequency-based analysis of resting-state fMRI signals revealed that functional connectivity between the right frontoparietal network and primary and dorsal occipital networks is linked to visual processing speed. This link was present in the slowest, typical frequency of the fMRI signal but also in the higher frequencies that are routinely discarded. These findings imply that the coordinated spontaneous activity between right frontoparietal and occipital regions supports the individual potential of the visual system for efficient processing.

## Introduction

Visual information processing requires an efficient visual attention system. The speed of information uptake in a given unit of time can be estimated using the mathematical framework of the ‘theory of visual attention’ (TVA; Bundesen, 1990). Within the TVA framework, visual processing speed (VPS) is estimated based on an individual’s accuracy in a whole-report task requiring the observer to report as many letters as possible from briefly presented displays. The estimated VPS represents a latent-level parameter that is relatively constant across diverse conditions (e.g., Finke et al., 2005). At a neural level, the computations that determine VPS, i.e., the selection of visual features, involve frontal, parietal, and limbic (control) areas in conjunction with occipital (visual-coding) areas (Bundesen et al., 2005). A prior resting-state functional magnetic resonance imaging (rs-fMRI) study showed that VPS relates to the inter-network functional connectivity (inter-FC) between the cingulo-opercular (COn) and the right frontoparietal (RFPn) networks (Ruiz-Rizzo et al., 2018)— both of which comprise frontal, parietal, and limbic areas. However, empirical evidence for the link between VPS and the inter-FC of COn and RFPn with occipital areas is missing. The present study set out to provide this evidence.

COn regions, such as the prefrontal, insular, and midcingulate cortices (Seeley et al., 2007; Dosenbach et al., 2008), increase their functional connectivity with the occipital cortex during eyes open versus eyes closed (Riedl et al., 2016). These regions exhibit sustained activity in tasks involving active visual processing (Sestieri et al., 2013). Similarly, the volume of the white matter tracts underlying RFPn regions (i.e., dorsolateral prefrontal cortex and areas around the intraparietal sulcus; Dosenbach et al., 2007) predict faster stimulus detection in visuospatial attention tasks (Thiebaut de Schotten et al., 2011), and more right-hemispheric lateralization of the inferior fronto-occipital fasciculus has been associated with higher values of the TVA-based VPS parameter (Chechlacz et al., 2015). The present study adds to the extant literature by shedding light on how inter-FC of COn and RFPn with occipital regions support VPS. Rs-fMRI allows measuring the correlation between brain regions’ spontaneous (i.e., intrinsic) hemodynamic activity (Raichle, 2011). We, thus, used rs-fMRI to study the intrinsic inter-FC of COn and RFPn with visual networks. Rs-fMRI data are typically temporally filtered (0.01-0.1 Hz) to remove non-neural scanner signal drifts (low frequencies) and cardio-respiratory signals (high frequencies) (Cordes et al., 2001). However, the spectral centroid (or ‘center of gravity’ of the frequencies) of COn, RFPn, and visual networks lies around the *upper* limit of the traditionally filtered frequency range (i.e., 0.098, 0.090, and 0.090-0.118 Hz, respectively; Ries et al., 2018). Further, *within*-network functional connectivity can be stronger in the typical range (< 0.08 Hz) for dorsal prefrontal regions (RFPn), but above 0.08 Hz for insular and orbitofrontal areas (COn, Salvador et al., 2008).

Previous rs-fMRI studies (e.g., Zuo et al., 2010; Gohel and Biswal, 2015; Wang et al., 2018) have examined all frequencies in their signal by adopting the slowest, supra-second oscillatory ranges derived from electrophysiological measures of neuronal activity (Penttonen and Buzsáki, 2003). This approach has revealed a similar spatial extent of functional connectivity *within* COn, RFPn, and visual networks across frequencies, with most of their total power in intermediate ranges (i.e., 0.073 to 0.198 Hz; Gohel and Biswal, 2015). Following this approach, here we used the entire frequency spectrum available in our rs-fMRI data (i.e., 0.01 to 0.4 Hz) and analyzed discrete frequency bins, including one ‘reference’ bin (i.e., entire spectrum). We expected significant inter-FC between COn, RFPn, and visual networks across frequency bins (Gohel and Biswal, 2015; Wang et al., 2018). We furthermore expected the inter-FC of COn and RFPn (respectively) with visual networks to be significantly linked to VPS and explored whether this link is frequency-specific.

## Methods

### Participants

Forty-eight healthy adults (age range: 20-50 years; 27 females), taken from the ‘INDIREA’ Munich cohort published in Ruiz-Rizzo et al. (2019), were included in the current study (see demographic information in Table 1), although the current hypotheses and analyses were generated independently. The study was approved by the ethics committee of the Faculty of Psychology and Educational Sciences of LMU Munich, and all participants provided written informed consent. To avoid strong age effects (e.g., on the VPS parameter; McAvinue et al., 2012), we selected all participants younger than 50 years from the original cohort. All of them exhibited normal psychomotor speed performance in a neuropsychological paper-and-pencil task (TMT-A, Reitan, 1958; Tombaugh, 2004), all had normal or corrected-to-normal visual acuity, and none was suffering from psychological or neurological disorders potentially affecting cognition, or from diabetes or color-blindness. To further ensure health status, participants additionally completed demographic and behavioral (e.g., Beck Depression Inventory, BDI; Beck et al., 1996) self-report questionnaires, as well as a test of crystallized intelligence (i.e., the multiple-choice vocabulary test: “Mehrfachwahl-Wortschatz-Test”; Lehrl et al., 1999). Thus, we assumed the relationship between inter-FC and our TVA-based measure of VPS to be uncontaminated by potential pathological influences.

**Table 1.**
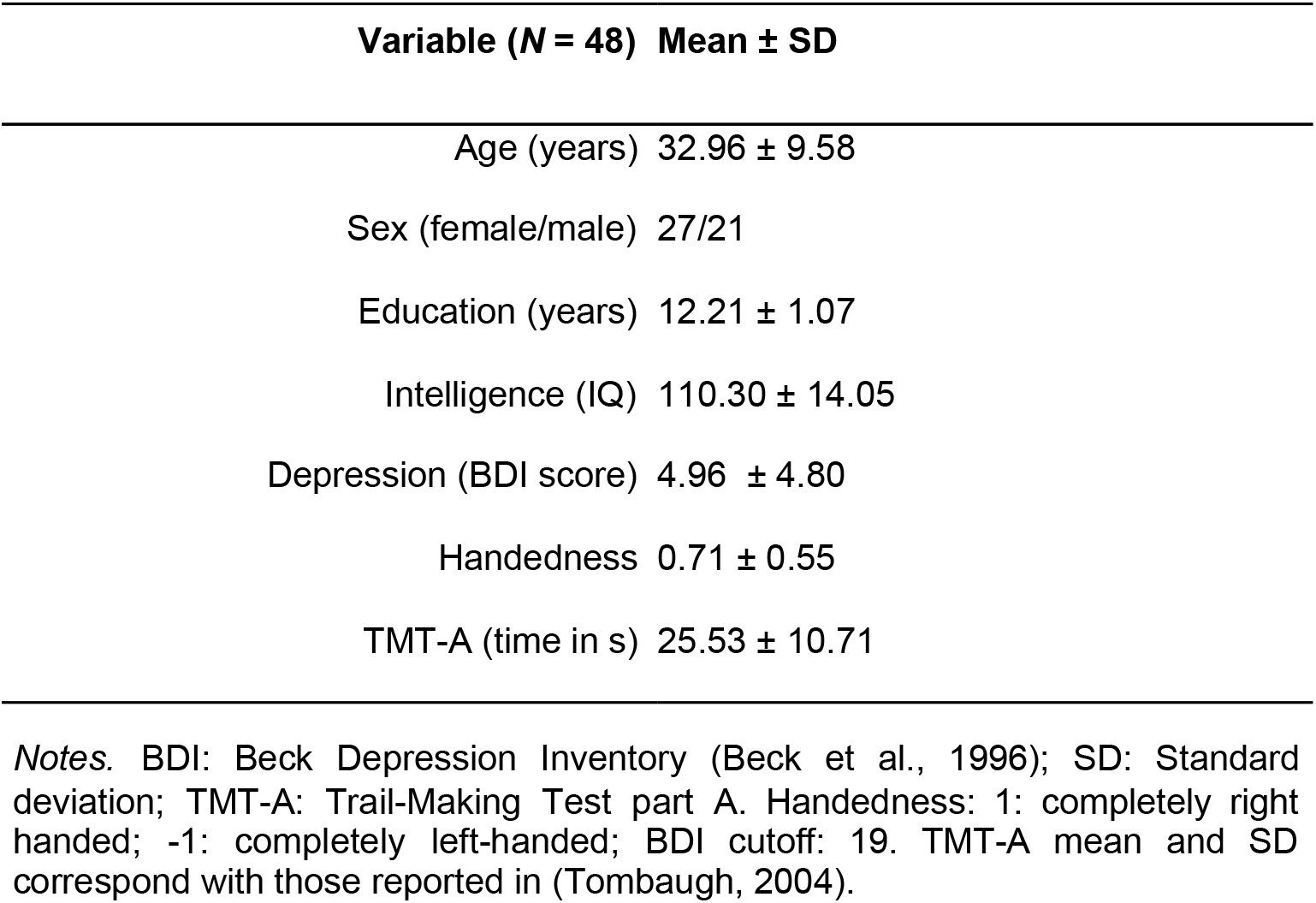
Demographic variables

Details on data acquisition are reported elsewhere (Ruiz-Rizzo et al., 2019). Briefly, after screening for inclusion and exclusion criteria, participants underwent rs-fMRI at the Klinikum rechts der Isar (Munich, Germany) in a first session. In a second session, participants performed the TVA-based whole-report task.

### Experimental Design and Statistical Analyses

#### Assessment and estimation of VPS parameter *C*

A whole-report task based on TVA (Bundesen, 1990) was used to estimate the TVA VPS parameter or parameter ‘*C*’. In this task, arrays of four letters were presented in an imaginary semicircle under different exposure durations determined in a pretest before the actual task. To identify individual shortest exposure durations, we used a staircase procedure in the pretest, which included four blocks of 12 trials each: four ‘adjustment’ trials, four trials with unmasked displays presented for 200 ms, and four masked displays presented for 250 ms. Each block started with one (of four) adjustment trial displayed for 80 ms, immediately followed by a post-display mask (see below). If the participant reported at least one letter correctly, the exposure duration was decreased by 10 ms in the following three adjustment trials within each block, until the participant could no longer report one letter correctly (i.e., the shortest exposure duration). If this point was reached before the last of the 16 adjustment trials, the exposure duration was held constant for the remaining. Setting the exposure duration that short ensured obtaining a valid estimate of the visual perceptual threshold parameter. Then, based on the shortest exposure duration, longer values were added to obtain report performance across the whole range from near-floor to near-ceiling and thus allow for a more precise TVA-based parameter estimation. In the actual whole-report task, on each trial, red letters were presented on either the left or the right side (counterbalanced) of a fixation point located in the screen center (Figure 1). Four blue items (shapes made of letter parts) were presented on the corresponding opposite site to balance visual stimulation. Letter stimuli were randomly chosen from the set (A, B, D, E, F, G, H, J, K, L, M, N, O, P, R, S, T, V, X). In a given trial, a particular letter appeared only once, and each letter was equally frequent within a block. Participants were instructed to report, verbally, all letters they were “fairly certain” they had seen (i.e., to avoid too conservative or too liberal a report criterion). Only report accuracy, but not report order or speed, was considered to assess performance.

**Figure 1.**
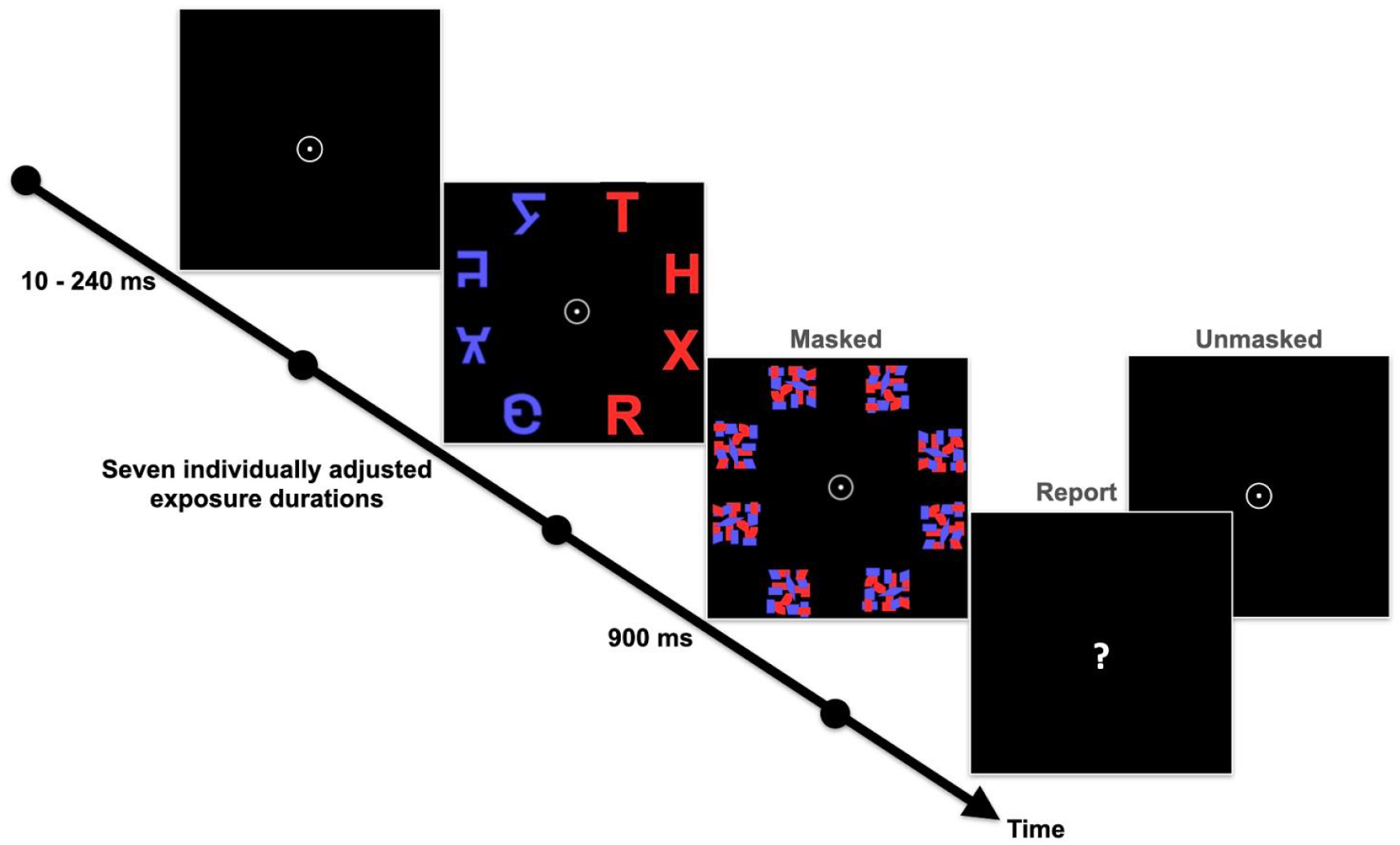
Whole-report task used to estimate the visual processing speed parameter *C*. Participants are asked to report all letters they are fairly certain they have seen, placing emphasis on accuracy, and not speed, of verbal report.

The task consisted of 400 trials, split into ten blocks of 40 trials each. Within a block, in fifteen of the trials, masks immediately followed stimuli presentation. These masks were jumbled blue and red squares, presented for 900 ms at each stimulus location to counteract visual persistence (Figure 1). Masked trials were presented for five different exposure durations (three times each). The use of varying exposure durations set for each individual was intended to increase precision in the TVA-based parameter estimation by allowing variability in report performance. The remaining 25 block trials were unmasked to add a component of iconic memory buffering (Sperling, 1960) to the estimation and ensure a valid and reliable TVA parameter fitting. These unmasked trials were presented for either the second shortest masked duration (three trials) or 200 ms (22 trials)—We chose the duration of 200 ms because electroencephalographic measures were simultaneously obtained to analyze event-related potentials (not reported in this study). Furthermore, as the shortest trial was too brief for visual perception, the second shortest was presented. Hence, in each block, trials were presented for seven *effective* exposure durations (five masked and two unmasked).

The VPS parameter, *C*, was estimated by modeling participants’ report accuracy as a function of the effective exposure duration, using a maximum likelihood-fitting algorithm (Bundesen, 1990; Kyllingsbæk, 2006; Dyrholm et al., 2011). Specifically, performance was modeled by an exponential growth function characterizing the increase in the probability of correct letter report with increasing effective exposure duration. The VPS parameter *C* represents the rate of uptake of visual information (in numbers of elements per second) and is given by the function’s slope at its origin. Although not in the focus of the present study, three additional parameters were estimated, namely, parameter *t*_*0*_, parameter *K*, and parameter *μ*. Parameter *t*_*0*_, the function’s origin, indicates the longest exposure duration (in ms) below which information uptake is effectively zero and represents the visual perceptual threshold. Parameter *K*, the function’s asymptote, indicates the maximum number of elements that can be simultaneously encoded in the visual short-term memory store. Parameter *μ* reflects the duration of iconic memory buffering in unmasked trials.

#### Statistical analyses

To statistically compare networks’ inter-FC *Z*-values across frequency bins, we performed repeated-measures analyses of variance (ANOVA), one for each relevant network pair, with frequency bins as the within-subject factor and *Z*-values for a particular network pair as the dependent variable. The relevant network pairs were COn and visual networks, RFPn and visual networks, COn and RFPn, and all visual networks. After each repeated-measures ANOVA, *post-hoc* tests were performed on individual *Z*-values for the respective network pairs. When Mauchly’s test indicated that the assumption of sphericity had been violated, Greenhouse-Geisser correction was applied. Bonferroni correction (*p*_*0.05/6*_ = 0.008 for comparisons involving COn, *p*_*0.05/3*_ = 0.017 for the rest of networks) was applied to *post-hoc* paired-sample *t*-tests. These analyses were performed in R 4.0.0 (R Core Team, 2020; https://www.R-project.org/; R.Studio v. 1.2.5042; RStudio Team, 2020; https://www.rstudio.com/).

We analyzed the relationship between inter-FC of COn and RFPn with visual networks and VPS parameter *C* by multiple regressions. Specifically, individual values of *C* were predicted from inter-FC *Z*-values in each frequency bin (Slow-5, Slow-4, Slow-3, Slow-2), frequencies altogether (henceforth named “Global”; see *Temporal filtering of rs-fMRI data* below), age (as after exclusion of high age the range was still 30 years), and framewise displacement (a measure of head movement in the scanner from frame to frame). Five multiple regression models were tested, one for each visual network’s inter-FC with COn and RFPn. Therefore, each model consisted of 13 predictors: six for the inter-FC of COn with a particular visual network across frequency bins and Global; five for the inter-FC of RFPn with the same visual network across frequency bins and Global; one for age; and one for framewise displacement. Note that the predictors involving COn were six instead of five because there were two subcomponents of COn in Slow-3 instead of one (see *Brain network selection* in the Results section). The goal of this analysis was to determine whether any of the predictors (i.e., inter-FC between a visual network and COn and RFPn, across frequency bins) in any of the models (that differed by i.e., visual network) was associated with parameter *C* — as we had no priors about one specific visual network or frequency bin being more relevant than the others. A second goal was to determine whether those associations were frequency-specific. Thus, we compared the beta coefficients in the models where significant predictors were found with linear *post-hoc* contrasts. Results were deemed significant if *p* < 0.05.

#### MRI data acquisition

Image acquisition was conducted on a Philips Ingenia 3T MRI-scanner (Philips Healthcare, Best, the Netherlands) with a standard 32-channel SENSE head coil, located in the Klinikum rechts der Isar, Munich (Germany). During the rs-fMRI session, participants were asked to try to avoid thinking about anything in particular, moving, or falling asleep. Head motion was restrained throughout the scanning session by foam padding around participants’ heads, and scanner noise was reduced by providing participants with earplugs and headphones. Six-hundred T2*-weighted blood oxygenation level-dependent (BOLD)-fMRI volumes were acquired per participant, using a multiband echo-planar imaging (EPI) sequence, with a 2-fold in-plane SENSE acceleration (SENSE factor, S = 2; Preibisch et al., 2015) and an M-factor of 2 (repetition time, 1250 ms; time to echo, 30 ms; flip angle, 70°; 40 slices; 3-mm slice thickness and 0.3-mm inter-slice gap; voxel size, 3 × 3 × 3.29 mm^3^; matrix size, 64 × 64). Anatomical detail was achieved by a higher resolution T1-weighted volume, acquired with a 3D magnetization prepared rapid acquisition gradient echo (MPRAGE) sequence (repetition time, 9 ms; time to echo, 4 ms; flip angle, 8°; 170 slices; voxel size, 1 mm^3^; matrix size, 240 × 240).

#### Rs-fMRI data preprocessing

As we were interested in examining frequencies that could also include physiological non-neural (e.g., respiratory) or scanner ‘noise’ signals (e.g., > 0.1 Hz), we reduced the possibility of including those non-neural signals by applying physiologic estimation by temporal independent component analysis (PESTICA; Beall and Lowe, 2007). PESTICA estimates the breathing and pulse signals from the data by performing slice-wise temporal ICA, identifying noise components per slice, and implementing signal correction (Beall and Lowe, 2007). We used this approach because no respiratory or cardiac signals were directly measured during rs-fMRI.

Next, we pre-processed the PESTICA-corrected rs-fMRI data using DPARSF (Data Processing Assistant for Resting-State fMRI; Yan and Zang, 2010), running on MATLAB (2016a, The Mathworks, Inc., Natick, U.S.A.). The preprocessing steps included discarding the first 5 volumes to remove initial T1 saturation; slice-timing correction; reorienting to anterior commissure-posterior commissure plane; realignment; co-registration to the high-resolution, structural image; DARTEL (Diffeomorphic Anatomical Registration using Exponentiated Lie algebra; Ashburner, 2007) segmentation into the three tissue types (gray matter, white matter, and cerebrospinal fluid); normalization to the Montreal Neurological Institute (MNI) space; spatial smoothing with a 4 mm full-width-at-half-maximum Gaussian kernel, and detrending. Moreover, the signals from white matter and cerebrospinal fluid, the six head motion parameters and their corresponding first derivatives, and the ‘bad’ frames (based on the framewise displacement metric of Power et al., 2012, as implemented in DPARSF) were regressed out from the rs-fMRI data. The global signal was not regressed out during preprocessing to avoid possible spurious anticorrelations.

#### Temporal filtering of rs-fMRI data

Following the frequency ranges of the slowest oscillations as defined in Penttonen and Buzsáki (2003) and on previous rs-fMRI studies (e.g., Zuo et al., 2010; Gohel and Biswal, 2015), we focused on four frequency bins: ‘Slow-5’ (0.01-0.027 Hz), ‘Slow-4’ (0.027-0.073 Hz), ‘Slow-3’ (0.073-0.198 Hz), and ‘Slow-2’ (0.198-0.4 Hz). The upper limit of Slow-2, and our highest frequency possible, was defined based on the Nyquist frequency for our 1.25-s repetition time, i.e., (1/1.25)/2 Hz. The current sampling frequency did not permit including the actual upper limit of Slow-2 (i.e., 0.5 Hz), and ‘Slow-1’ (0.5–0.75 Hz) (*cf.* Gohel and Biswal, 2015). We used these bins to temporally filter the preprocessed data. Thus, we obtained five data versions, each with a different frequency composition, namely, the four frequency bins and one ‘Global,’ encompassing all bins (i.e., 0.01-0.4 Hz), for comparison. Note that these versions solely differed in their spectral content and that Global was included as a reference, to better appreciate the possible frequency specificity of inter-FC and its association with VPS.

#### Independent component analysis (ICA) and dual regression

Spatial group ICA was used to decompose the preprocessed BOLD-fMRI volumes into 75 independent components (ICs; following Allen et al., 2011) with FSL (v. 5.0.7; Jenkinson et al., 2012) MELODIC (v. 3.14; Smith et al., 2004). Dual regression was performed on all ICs to generate participant-specific time courses (stage 1) and spatial maps (stage 2) (Beckmann et al., 2009). The participant-specific time courses were used as input for the inter-FC analysis. Both ICA and dual regression were performed separately for each of the four frequency bins and for the Global data.

#### Brain network selection

For each frequency bin, we obtained the ICs’ spatial cross-correlation values with resting-state network templates (Allen et al., 2011) using FSL’s *fslcc* command (https://surfer.nmr.mgh.harvard.edu/pub/dist/freesurfer/tutorial_packages/OSX/fsl_501/src/avwutils/fslcc.cc). As we were interested in networks relevant for VPS, we selected the two ICs, as defined in Allen et al. (2011), that showed the highest correlation coefficients with, correspondingly, COn (Allen et al.’s Salience, IC 55), RFPn (Allen et al.’s IC 60), and visual networks (Allen et al.’s IC 39, IC 46, IC 48, IC 59, IC 64, and IC 67). Next, the two ICs that correlated the highest were visually inspected and the one including the most relevant regions of each network were chosen. These regions included the anterior insula and anterior midcingulate cortex for COn; the right middle frontal gyrus and anterior inferior parietal lobule for RFPn; and striate and extrastriate cortex, as well as lateral geniculate nucleus of the thalamus, for the visual networks (Uddin et al., 2019). After visual inspection, if both ICs included parts of the relevant regions in a complementary manner (i.e., the network was split), both were selected. If both ICs included the relevant regions but these were unidentifiable from the overall spatial pattern of the IC (i.e., the ICs additionally included other brain regions or noise patterns), none was selected.

#### Inter-FC analyses

To calculate the individual inter-FC of network pairs, the specific time courses (i.e., those derived from stage 1 of the dual regression) of the networks of interest were correlated for each participant. The resulting *r-*value matrices were Fisher *Z*-transformed and averaged across participants to obtain a group-level inter-FC matrix. One-sample *t*-tests were computed on the group mean Fisher-*Z*-transformed correlation matrix and the false discovery rate method (FDR; Benjamini and Hochberg, 1995) was used to correct for multiple comparisons (*p* < 0.05 and *q* < 0.05). We repeated this step for each frequency bin (including Global). This analysis was performed with custom code written in MATLAB (Ruiz-Rizzo and Küchenhoff, 2020).

#### Code Accessibility

Analysis scripts for the inter-FC and the statistical analyses and analyzed neuroimaging data are publicly available and can be accessed at (https://osf.io/nhqg3/). Analyzed behavioral data will also be accessible to qualified researchers upon request.

## Results

### Brain network selection

Coefficients for the cross-correlation of ICs with the network templates of Allen et al. (2011) are listed in Table 2. COn was consistently identified across all frequency bins and Global. For Slow-3, two subcomponents of COn were identified, one centered on the insula and the other on the anterior cingulate cortex (ACC) (Figure 2A, top row, third column; IC32 and IC42 respectively). Thus, both ICs (i.e., IC32 and IC42) were included in further analyses. RFPn was consistently identified across all frequency bins and Global (Figure 2A, bottom row). Finally, five (out of six) of Allen et al.’s visual system networks (named here as Vis-39, Vis-46, Vis-59, Vis-64, and Vis-67 following Allen et al’s IC numbering) were consistently identified across all frequency bins and Global (Figure 2B). Allen et al.’s IC 48 could be identified only in Global and Slow-4 and, thus, was excluded from further analyses.

**Table 2.** ICs with highest correlation coefficients with network templates across frequency bins (and Global, for comparison)

| Network<br>template | Frequency bin |  |  |  |  |
| --- | --- | --- | --- | --- | --- |
|  | Slow-5 | Slow-4 | Slow-3 | Slow-2 | Global |
| COn | **IC20 (0.63)** | IC32 (0.52) | **IC32 (0.38)** | **IC51 (0.27)** | IC27 (0.39) |
|  | IC15 (0.24) | **IC35 (0.41)** | **IC42 (0.31)** | IC23 (0.24) | **IC44 (0.36)** |
| RFPn | **IC7 (0.64)** | **IC4 (0.70)** | **IC2 (0.72)** | **IC61 (0.43)** | **IC3 (0.69)** |
|  | IC14 (0.25) | IC18 (0.21) | IC33 (0.27) | IC58 (0.26) | IC15 (0.33) |
| Vis-39 | **IC11 (0.49)** | **IC20 (0.43)** | **IC3 (0.44)** | IC52 (0.28) | **IC4 (0.49)** |
|  | IC14 (0.28) | IC40 (0.43) | IC33 (0.35) | **IC40 (0.22)** | IC15 (0.28) |
| Vis-46 | **IC2 (0.49)** | **IC8 (0.58)** | IC5 (0.47) | **IC11 (0.47)** | IC5 (0.46) |
|  | IC1 (0.35) | IC2 (0.36) | **IC10 (0.39)** | IC10 (0.31) | **IC11 (0.44)** |
| Vis-59 | **IC1 (0.51)** | **IC6 (0.47)** | **IC4 (0.54)** | **IC2 (0.51)** | **IC1 (0.52)** |
|  | IC32 (0.32) | IC17 (0.39) | IC63 (0.29) | IC3 (0.25) | IC28 (0.33) |
| Vis-64 | **IC5 (0.55)** | **IC1 (0.64)** | **IC5 (0.51)** | IC8 (0.49) | **IC5 (0.48)** |
|  | IC2 (0.44) | IC6 (0.35) | IC4 (0.41) | **IC10 (0.37)** | IC14 (0.45) |
| Vis-67 | **IC18 (0.47)** | **IC2 (0.48)** | **IC40 (0.48)** | IC44 (0.44) | IC38 (0.44) |
|  | IC43 (0.33) | IC39 (0.36) | IC10 (0.30) | **IC8 (0.31)** | **IC14 (0.38)** |

*Note.* Correlation coefficients between Allen et al. (2011)'s templates and the two independent components (ICs) with the highest values are shown separately for networks of interest and frequency bins. Selected ICs (based on correlation coefficients *and* visual inspection) are marked in bold. Allen et al.'s Salience (IC 55) and Right frontoparietal network (IC 60) correspond to the cingulo-opercular (COn) and RFPn, respectively, of the present study. Slow-5: 0.01-0.027 Hz; Slow-4: 0.027-0.073 Hz; Slow-3: 0.073-0.198 Hz; Slow-2: 0.198-0.4 Hz. Vis: visual networks.

| Network template | Frequency bin |  |  |  |  |
| --- | --- | --- | --- | --- | --- |
|  | Slow-5 | Slow-4 | Slow-3 | Slow-2 | Global |
| COn | **IC20 (0.63)** | IC32 (0.52) | **IC32 (0.38)** | **IC51 (0.27)** | IC27 (0.39) |
|  | IC15 (0.24) | **IC35 (0.41)** | **IC42 (0.31)** | IC23 (0.24) | **IC44 (0.36)** |
| RFPn | **IC7 (0.64)** | **IC4 (0.70)** | **IC2 (0.72)** | **IC61 (0.43)** | **IC3 (0.69)** |
|  | IC14 (0.25) | IC18 (0.21) | IC33 (0.27) | IC58 (0.26) | IC15 (0.33) |
| Vis-39 | **IC11 (0.49)** | **IC20 (0.43)** | **IC3 (0.44)** | IC52 (0.28) | **IC4 (0.49)** |
|  | IC14 (0.28) | IC40 (0.43) | IC33 (0.35) | **IC40 (0.22)** | IC15 (0.28) |
| Vis-46 | **IC2 (0.49)** | **IC8 (0.58)** | IC5 (0.47) | **IC11 (0.47)** | IC5 (0.46) |
|  | IC1 (0.35) | IC2 (0.36) | **IC10 (0.39)** | IC10 (0.31) | **IC11 (0.44)** |
| Vis-59 | **IC1 (0.51)** | **IC6 (0.47)** | **IC4 (0.54)** | **IC2 (0.51)** | **IC1 (0.52)** |
|  | IC32 (0.32) | IC17 (0.39) | IC63 (0.29) | IC3 (0.25) | IC28 (0.33) |
| Vis-64 | **IC5 (0.55)** | **IC1 (0.64)** | **IC5 (0.51)** | IC8 (0.49) | **IC5 (0.48)** |
|  | IC2 (0.44) | IC6 (0.35) | IC4 (0.41) | **IC10 (0.37)** | IC14 (0.45) |
| Vis-67 | **IC18 (0.47)** | **IC2 (0.48)** | **IC40 (0.48)** | IC44 (0.44) | IC38 (0.44) |
|  | IC43 (0.33) | IC39 (0.36) | IC10 (0.30) | **IC8 (0.31)** | **IC14 (0.38)** |

*Note.* Correlation coefficients between Allen et al. (2011)'s templates and the two independent components (ICs) with the highest values are shown separately for networks of interest and frequency bins. Selected ICs (based on correlation coefficients *and* visual inspection) are marked in bold. Allen et al.'s Saliency (IC 55) and Right frontoparietal network (IC 60) correspond to the cingulo-opercular (COn) and RFPn, respectively, of the present study. Slow-5: 0.01-0.027 Hz; Slow-4: 0.027-0.073 Hz; Slow-3: 0.073-0.198 Hz; Slow-2: 0.198-0.4 Hz. Vis: visual networks.

**Figure 2.**
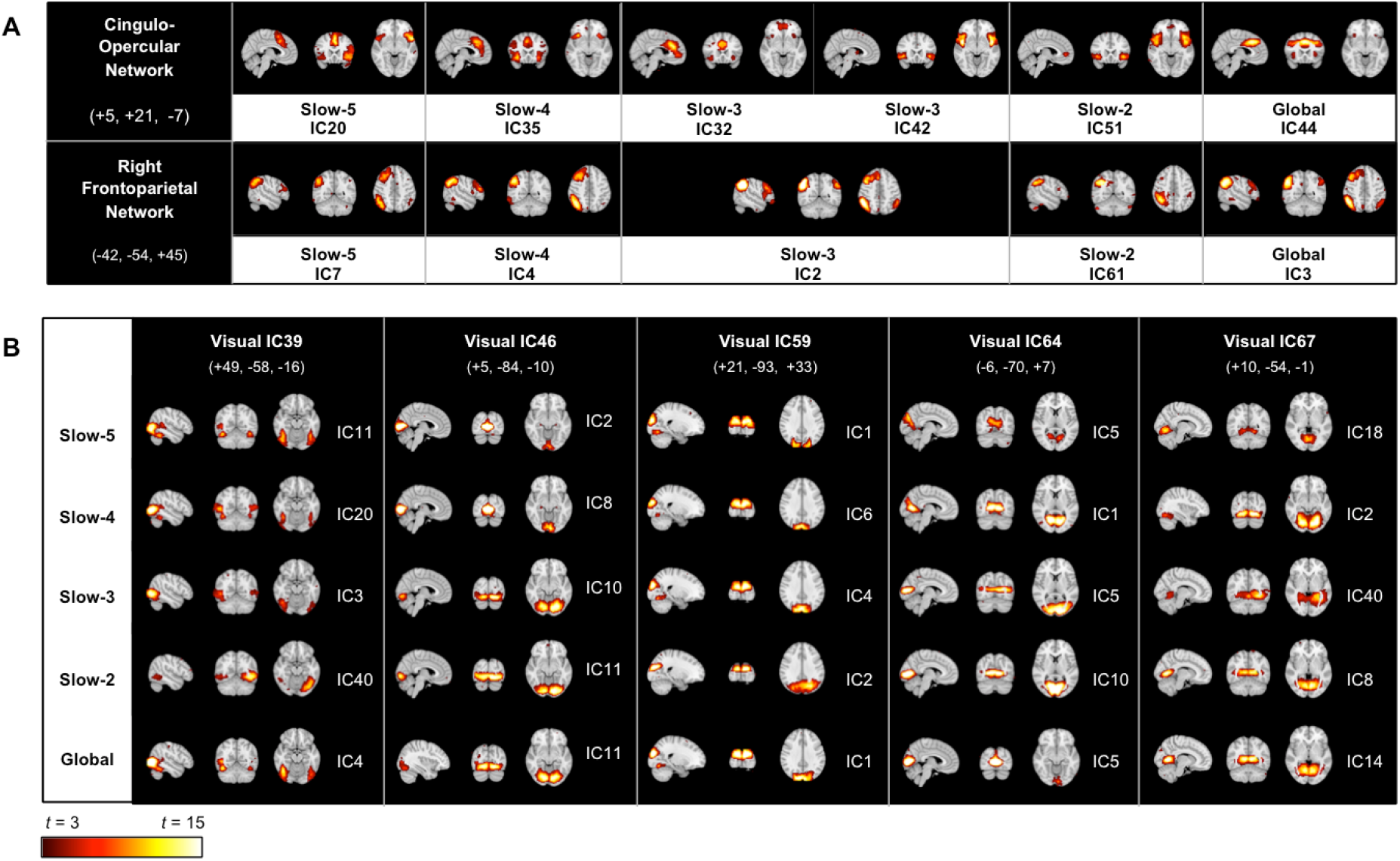
Networks of interest identified for each frequency bin. Independent components (IC) representative of the cingulo-opercular and right frontoparietal networks (A) and visual networks (B) (following Allen et al., 2011) in each frequency bin (Slow-5 to Slow-2) and Global, for comparison. Montreal Neurological Institute coordinates (x, y, z, in mm) correspond to the slices shown for each network. Slow-5: 0.01-0.027 Hz; Slow-4: 0.027-0.073 Hz; Slow-3: 0.073-0.198 Hz; Slow-2: 0.198-0.4 Hz.

### Inter-FC between COn, RFPn, and visual networks

The group average correlation matrix, per frequency bin, between COn, RFPn, and visual networks is depicted in Figure 3, and the corresponding statistical comparison across frequency bins, for COn, RFPn, and visual network pairs is shown in Figure 4. COn exhibited significant inter-FC with three visual networks (Vis-39, Vis-59, and Vis-67) in three frequency bins: Slow-4, Slow-3, and Slow-2. The inter-FC with Vis-39 was negative in both Slow-4 (*Z* = -0.28, *p* < .001) and Slow-2 (*Z* = -0.09, *p* = 0.004). The inter-FC with Vis-59 was negative in Slow-4 (*Z* = -0.16, *p* < 0.001) and positive in Slow-3 (ACC-subcomponent: *Z* = 0.08, *p* = 0.011; insula-subcomponent: *Z* = 0.14, *p* < 0.001). Finally, the inter-FC with Vis-67 was also negative in Slow-4 (*Z* = -0.12, *p* = 0.004) and positive in Slow-3 (insula-subcomponent: *Z* = 0.12, *p* < 0.001). As a reference, when all frequencies were considered together (i.e., Global), COn exhibited significant negative inter-FC with Vis-39 (*Z* = -0.12, *p* < 0.0001) and Vis-59 (*Z* = -0.10, *p* < 0.0001) and positive with Vis-46 (*Z* = 0.07, *p* = 0.004). When comparing across frequency bins (Figure 4A), we found the magnitude of the inter-FC between COn and visual networks to differ significantly [*F*(3.38, 812.6) = 33.67, *p* < 0.001, *η*^*2*^ = 0.09]. In particular, the (positive) inter-FC between COn’s insula-subcomponent and visual networks in Slow-3 was significantly stronger compared to the inter-FC between (single component) COn and visual networks in all other frequency bins (all Bonferroni-corrected *p*-values < 0.001).

**Figure 3.**
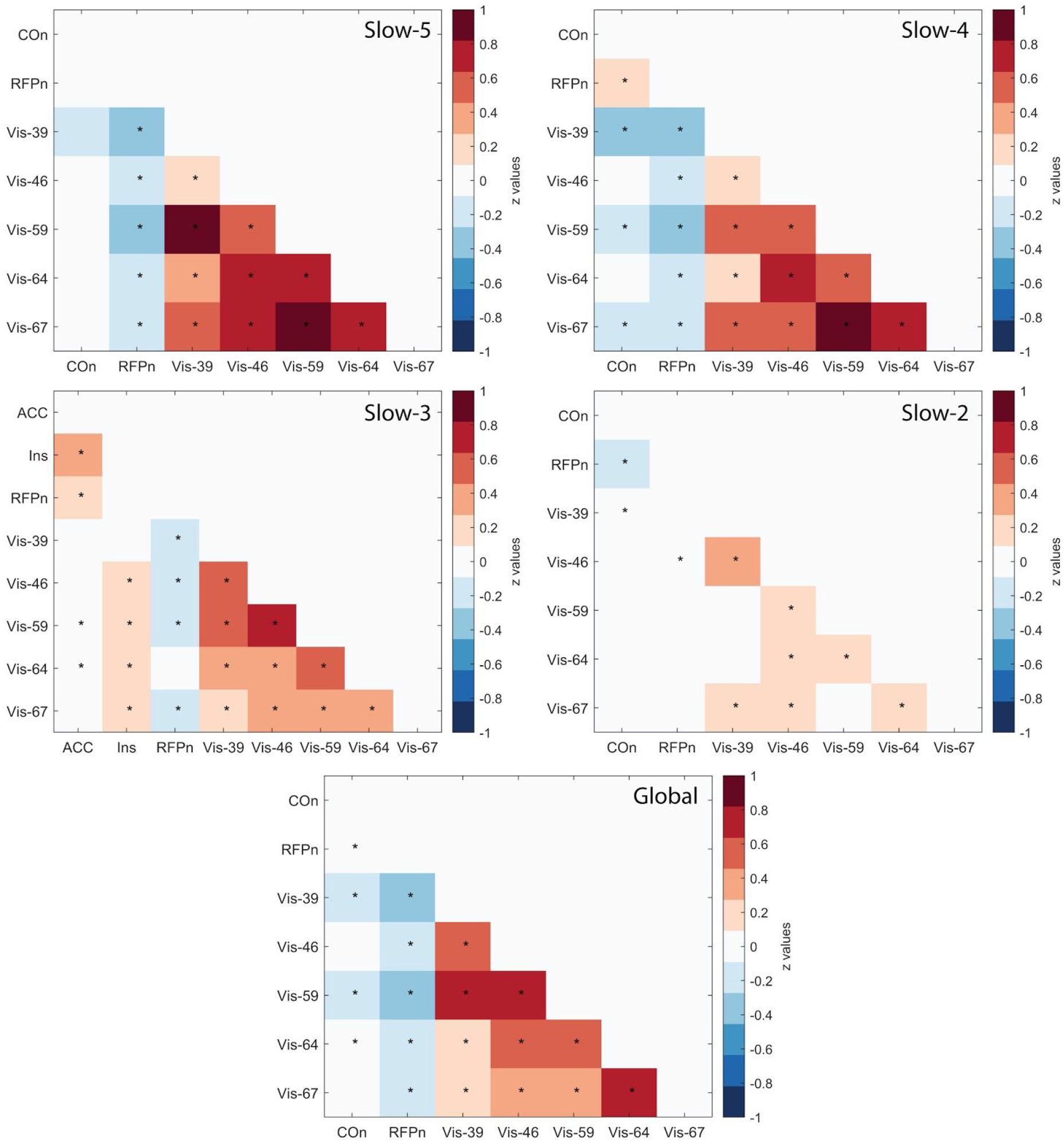
Group average correlation matrices between visual processing speed (VPS) relevant networks. Correlation matrices representing functional connectivity (*Z*-values) between higher-order networks previously shown relevant for VPS (cingulo-opercular network, COn, and right frontoparietal network, RFPn) and visual networks (Vis) across frequency bins (Slow-5 to Slow-2) and Global (frequencies altogether), for reference. Red indicates positive correlations, whereas blue and white indicate, respectively, negative and around-zero correlations. FDR-corrected (*q* < 0.05) significant values are marked with a ‘*’. See text for specific frequency ranges included in each bin.

**Figure 4.**
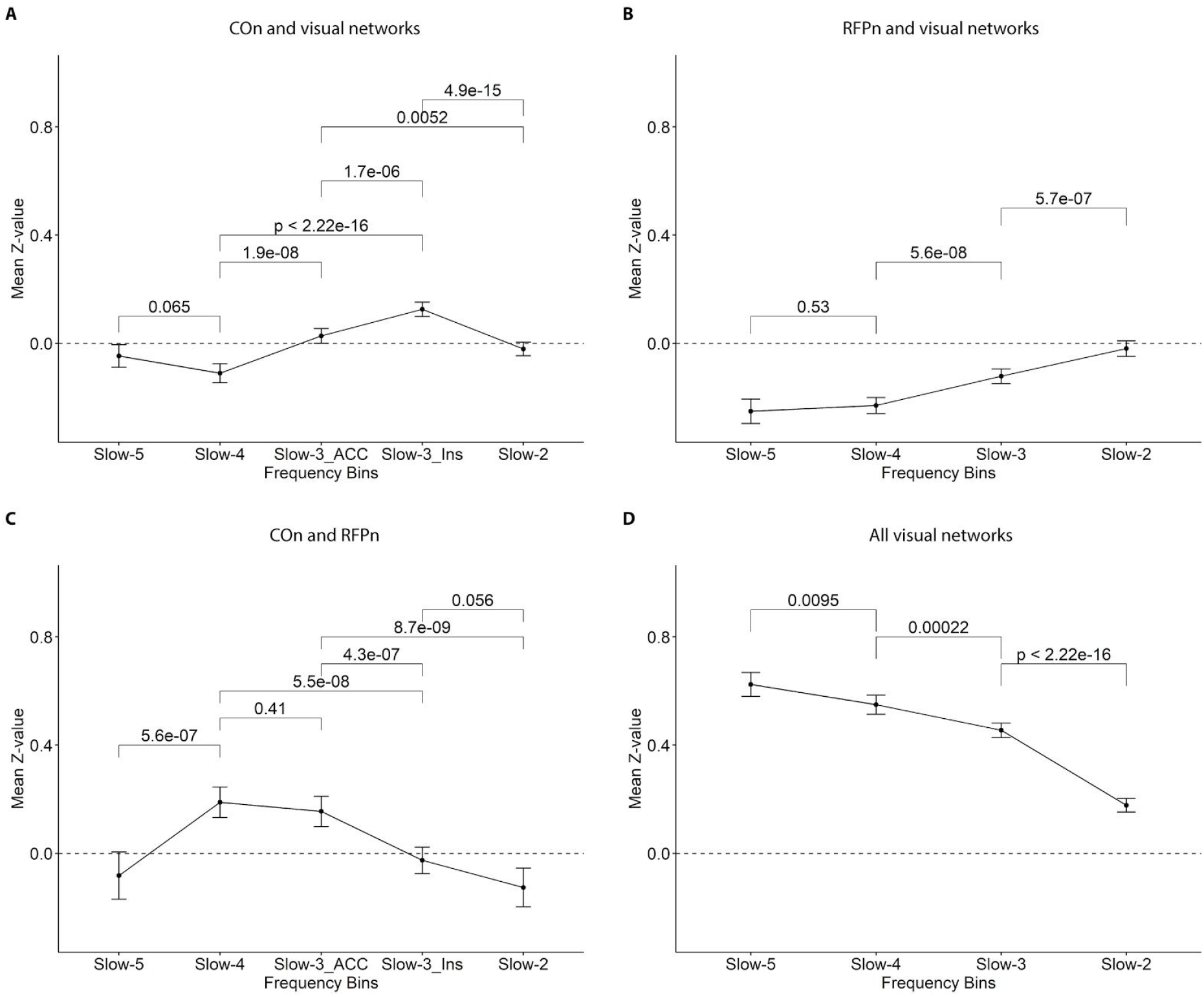
Mean inter-FC between cingulo-opercular, right frontoparietal, and visual networks across frequency bins. (A) The mean *Z*-value of the inter-FC between the cingulo-opercular (COn) and all visual networks was positive and strongest in Slow-3 with the COn’s insula-subcomponent (Slow3_Ins). (B) The mean *Z*-value of the inter-FC between the right frontoparietal network (RFPn) and all visual networks was negative and strongest in Slow-5 and Slow-4. (C) The mean *Z*-value of the inter-FC between COn and RFPn was positive and strongest in Slow-4 and Slow-3 with the COn’s ACC-subcomponent (Slow-3_ACC). (D) The mean *Z*-value of the inter-FC among visual networks was positive and strongest in Slow-5. Significance was determined at Bonferroni-corrected *p*_*0.05/6*_ = 0.008 (for A and C, where COn was subdivided into two subcomponents) and *p*_*0.05/3*_ = 0.017 (for B and D). Bars represent 95% confidence intervals.

The RFPn exhibited significant negative inter-FC with the same three visual networks as COn (Vis-39, Vis-59, and Vis-67) in all frequency bins (Figure 3; e.g., Vis-59: *Z* = -0.38 in Slow-5; Vis-39: *Z* = -0.35 in Slow-4; Vis-67: *Z* = -0.10 in Slow-3; all *p*-values < 0.001) but Slow-2, in which the only significant inter-FC was observed with Vis-46 (*Z* = -0.07, *p* = 0.021). As a reference, in Global, RFPn showed significant negative inter-FC with all visual networks (*Z*-values’ range: -0.37 to -0.19; all *p*-values < 0.0001). When comparing across frequency bins (Figure 4B), we again found the inter-FC between RFPn and visual networks to differ significantly [*F*(2.28, 544.92) = 50.94, *p* < 0.001, *η*^2^ = 0.11]. In particular, the mean inter-FC was decreasingly less negative with increasing frequency, indicating that correlations were stronger and more negative in slower than in faster frequency bins. Bonferroni-corrected *post-hoc* tests revealed that inter-FC between RFPn and visual networks was equally strong in Slow-5 (mean *Z* = -0.25) and Slow-4 (mean *Z* = -0.23), but decreased in magnitude in Slow-3 (mean *Z* = -0.12, *p* < 0.0001, in comparison to all other bins) and Slow-2 (mean *Z* = -0.02, *p* < 0.001, in comparison to all other bins).

COn showed significant inter-FC with RFPn in the following frequency bins (Figure 3): positive in Slow-4 (*Z* = 0.19, *p* < 0.0001) and Slow-3 (ACC-subcomponent: *Z* = 0.15, *p* < 0.0001), and negative in Slow-2 (Z = -0.13, *p* < 0.001). In Slow-5, inter-FC with RFPn was non-significant (*Z* = -0.08, *p* = 0.065). As a reference, in Global, the inter-FC between these two networks was positive (*Z* = 0.08, *p* < 0.001). Comparison across frequency bins (Figure 4C) revealed the inter-FC between RFPn and COn to differ significantly [*F*(2.90, 136.49) = 18.30, *p* < 0.001, *η*^2^ = 0.24]. Bonferroni-corrected *post-hoc* tests showed that the inter-FC between RFPn and COn was significantly more positive in Slow-4 and Slow-3 (ACC-subcomponent only) than in the other frequency bins or the insula-subcomponent in Slow-3 (all *p*-values < 0.001; Figure 4C).

Finally, for all visual networks, there was positive inter-FC throughout the entire frequency range (range: *Z* = 0.84 in Slow-5 to 0.06 in Slow-2; Figure 3). As a reference, in Global, the inter-FC among all visual networks was significantly positive, too [*Z*-values’ range: -0.75 (Vis-59 and Vis-48) to -0.12 (Vis-39 and Vis-67)]. When comparing across frequency bins (Figure 4D), although always positive, the inter-FC between visual network pairs differed significantly [*F*(3.38, 812.6) = 33.67, *p* < 0.001, *η*^2^ = 0.09]. Specifically, inter-FC decreased in strength with increasing frequency (Figure 4D). For example, the inter-FC between Vis-39 and Vis-59 was of 0.83 (*p* < 0.0001) in Slow-5, 0.59 (*p* < 0.0001) in Slow-4, 0.54 (*p* < 0.0001) in Slow-3, but only of 0.06 in Slow-2 (*p* = 0.037, *n.s*. after FDR correction).

In summary, we found that the mean inter-FC of COn with visual networks was positive and strongest in Slow-3 (insula-subcomponent); the mean inter-FC of RFPn with visual networks was negative and strongest in Slow-5 and Slow-4; the mean inter-FC between COn and RFPn was positive and strongest in Slow-4 and Slow-3 (ACC-subcomponent); and the mean inter-FC between visual networks was positive and strongest in Slow-5. Next, we investigated whether there is an association of VPS *C* with the inter-FC between these networks and, further, whether this association is frequency-specific.

### Association of VPS parameter C with the inter-FC between COn, RFPn, and visual networks

Values for VPS parameter *C* ranged from 10.34 to 47.01 letters/s (mean = 22.90 ± 7.70). We conducted five linear regressions (one for each visual network: Vis-39, Vis-46, Vis-59, Vis-64, and Vis-67) to examine the associations between VPS parameter *C* values and the inter-FC of COn and RFPn with visual networks across frequency bins and Global, controlling for age and head motion.

Figure 5 depicts the predictors’ beta coefficients (and their corresponding 95% confidence intervals) of inter-FC of COn (A) and RFPn (B) with each visual network across frequency bins (depicted in different colors and geometrical shapes). Significant associations with parameter *C* were found only for the inter-FC of RFPn with two of the five visual networks (Vis-59 and Vis-64) (confidence intervals not including zero; Figure 5B). Specifically, more *negative* inter-FCs of RFPn with Vis-59 (Slow-5: β = -0.56, SE = 0.22, *p* = 0.014) and Vis-64 (Slow-5: β = -0.56, SE = 0.22, *p* = 0.015; Slow-2: β = -0.35, SE = 0.16, *p* = 0.034) were significantly associated with higher VPS parameter *C*. No further significant associations were observed (all *p*-values > 0.060).

**Figure 5.**
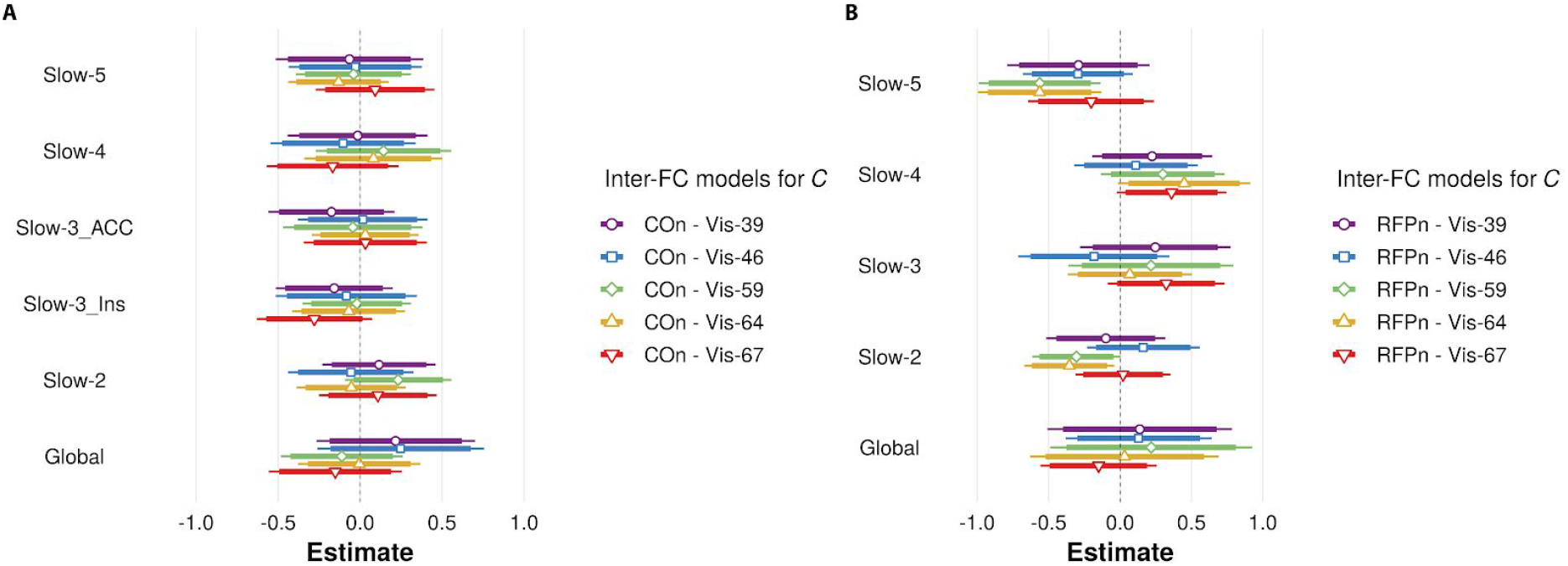
Estimates of linear regressions of visual processing speed (VPS) parameter *C* on inter-FC of cingulo-opercular network (COn) and right frontoparietal network (RFPn) with visual networks. The standardized coefficients (and their respective 95% confidence intervals) of five multiple regression models of VPS *C* on the inter-FC of COn (A) and the RFPn (B) with each visual network (Vis-), are depicted in five different colors and geometrical shapes. All models included as predictors the inter-FC of both COn and RFPn with one of the five visual networks, respectively (hence five regression models), across frequency bins and Global, age, and head motion. For clarity, inter-FC predictors are clustered on the *y*-axis and presented separately for COn (A) and RFPn (B). Estimates of age and head motion, as well as the of intercept, were omitted from the figure for simplicity. Significant estimates are those whose confidence interval does not cross the middle vertical (black dashed) line. Conventions for the regression models: Vis-39: purple, circle; Vis-46: blue, square; Vis-59: green, diamond; Vis-64: ocher, upward triangle; Vis-67: red, downward triangle.

To determine possible frequency specificity for the frequency bins that proved significant (i.e., inter-FC of RFPn and Vis-59 in Slow-5 and inter-FC of RFPn and Vis-64 in Slow-5 and Slow-2), we calculated *post-hoc* linear contrasts of the beta coefficients (i.e., Slow-5 > Slow-4; Slow-5 > Slow-3; Slow-5 > Slow-2; Slow-2 > Slow-4; and Slow-2 > Slow-3). In the regression model of inter-FC between Vis-59 and RFPn (green diamond in Figure 5B), contrasts indicated that the association was significantly different between Slow-5 and Slow-4 (*z* = -2.77, *p* = 0.016), marginally different between Slow-5 and Slow-3 (*z* = -2.24, *p* = 0.065), and not different between Slow-5 and Slow-2 (*p* = 0.685). Similarly, in the model of inter-FC of RFPn with Vis-64 (ocher upward triangle in Figure 5B), the association was significantly different between Slow-5 and Slow-4 (*z* = -3.43, *p* = 0.002) and Slow-2 and Slow-4 (*z* = -2.53, *p* = 0.022), but not between Slow-5 and Slow-3 (*p* = 0.102), Slow-5 and Slow-2 (*p* = 0.751), or Slow-2 and Slow-3 (*p* = 0.249). Thus, the association with VPS parameter *C* observed in Slow-5 for the inter-FC between RFPn and Vis-59 and Vis-64 appears to be also present (although less marked) in Slow-3 and Slow-2, arguing that it is not frequency-specific.

Surprisingly, the inter-FC of COn and visual networks was not significantly associated with VPS parameter *C* in any of the frequency bins and in any of the models (all *p*-values > 0.129; Figure 5A). We also found no significant results for the inter-FC of RFPn or COn with visual networks in Global in any of the models (all *p*-values > 0.158). None of the control variables showed significant associations with VPS parameter *C* in any of the models (age: all *p*-values > 0.255; head motion: all *p*-values > 0.400).

For completeness, we additionally examined the potential link between VPS parameter *C* and the inter-FC between COn and RFPn. We observed a significant association between the inter-FC of COn’s ACC-subcomponent and RFPn in Slow-3 only (β = -0.34, SE = 0.16, *p* = 0.043; *p*-values of all other frequency bins and Global > 0.269).

In summary, we found higher VPS parameter *C* to be significantly associated with a more negative inter-FC between RFPn and visual networks in Slow-5 (Vis-59 and Vis-64) and Slow-2 (Vis-64) only. However, contrary to our expectations, we did not find a significant association between VPS parameter *C* and the inter-FC of COn with visual networks for any frequency bin or Global.

## Discussion

In this study, we used theory of visual attention modeling and a frequency-based approach to examine whether the between-network functional connectivity (inter-FC) of the cingulo-opercular network (COn) and the right frontoparietal network (RFPn) with visual networks is associated with visual processing speed (VPS). We found that inter-FC of RFPn, but not that of COn, with visual networks was linked with VPS, and that this link was also observed beyond the typically analyzed frequency range. Filtering rs-fMRI data into four frequency bins (Slow-5: 0.01-0.027 Hz, Slow-4: 0.027-0.073 Hz, Slow-3: 0.073-0.198 Hz, and Slow-2: 0.198-0.4 Hz) revealed a functional subdivision for COn in Slow-3, with the strongest inter-FC with visual networks for COn’s insula-subcomponent and the strongest inter-FC with RFPn for COn’s anterior cingulate cortex (ACC)-subcomponent. Further, our approach revealed the strongest inter-FC between RFPn and visual networks in the slowest bins (Slow-5 and Slow-4). Our results are indicative of a frequency-specific inter-FC between higher-order and primary resting-state networks relevant for VPS and provide the first empirical evidence for a link between a latent VPS parameter and the intrinsic inter-FC between right frontoparietal (but not frontoinsular) and occipital regions.

### Inter-FC between RFPn and visual networks links with VPS

We found that the inter-FC between RFPn and visual networks, but not between COn and visual networks, was associated with the VPS parameter (Figure 5). This association was observed not only in the typically analyzed frequency range (Slow-5) but, notably, also in frequencies beyond this range (Slow-2 and, less strongly, Slow-3). The association with VPS was found for the inter-FC between RFPn and Vis-59 and Vis-64, i.e., networks that include, respectively, dorsal (adjoining parietal cortex) and primary visual areas (Allen et al., 2011). This finding is in accordance with well-established evidence that attentional control signals from (fronto)parietal regions of RFPn modulate sensory processing in visual cortices (Green and McDonald, 2008; Gilbert and Li, 2013; Scolari et al., 2015; Riedl et al., 2016) and that more right-hemispheric lateralization of the inferior fronto-occipital fasciculus associates with higher VPS (Chechlacz et al., 2015). In the present study, inter-FC was measured at rest and the VPS parameter *C* was obtained independently (i.e., from modeling accuracy in a whole-report task performed on a different day). Accordingly, this association would imply that the *intrinsic* inter-FC reflects the individual potential of the visual system to process information in an efficient manner, as the parameter *C* represents a latent-level measure of VPS (e.g., Finke et al., 2005).

Higher VPS parameter *C* was associated with *lower* (negative) inter-FC between RFPn and visual networks (Figure 3). Theoretically, VPS is determined by categorizations, i.e., selection of visual features (“object *x* has feature *i*”; Bundesen, 1990). In neural terms, such selection increases the firing rate of the cortical neurons coding for a particular feature (and, correspondingly, decreases the firing rate of neurons coding for other features) (Bundesen et al., 2005). Functional connectivity has been proposed to reflect fluctuations in cortical excitability (Raichle, 2011). Moreover, spontaneous, infra-slow neuronal fluctuations have been shown to underlie functional connectivity (Matsui et al., 2016). Thus, negative inter-FC between RFPn and visual networks suggests a neural implementation of the selection mechanism—a visual categorization would increase the activity of, specifically, the visual areas coding for a particular feature, but not of *all* visual areas (coding other features and where activity would decrease), thereby translating into a net decrease.

Contrary to what we expected, and unlike RFPn, the inter-FC between COn and visual networks was not associated with VPS parameter *C*. One possibility for this null finding may be that COn’s functional role in sustained cognitive control (see below) is not limited solely to visual processing and requires more interaction with higher-order networks (e.g., to control switching between default mode and central executive networks; Sridharan et al., 2008) than with unimodal networks. This possibility is further supported by our finding of an association of the inter-FC between COn and RFPn with VPS. COn and RFPn are two functionally well coupled but distinct networks associated with cognitive control (e.g., Dosenbach et al., 2007; Crittenden et al., 2016). Whereas COn has been suggested to be involved in cognitive control across longer periods of time, RFPn appears to adjust control more rapidly and dynamically (Dosenbach et al., 2008; Sadaghiani et al., 2012). The functional connectivity *within* COn and its inter-FC with RFPn have previously been shown relevant for the VPS parameter *C* (e.g., Ruiz-Rizzo et al., 2018). In light of the previous evidence, the current results suggest a possible hierarchical structure for the functional-connectivity-based correlates of the complex individual trait underlying VPS. This functional hierarchy would imply that COn might trigger sustained cognitive control, RFPn might then be stimulated to enhance the phasic response to incoming stimuli and thereby gate stimulus processing in visual (primary and dorsal occipital) regions. Assessing effective (i.e., directional) FC could provide further empirical support to this proposal.

### Inter-FC between VPS relevant networks is frequency-specific

Previous studies showed that resting-state networks can be observed at frequencies beyond the traditionally examined frequency range (0.01 - 0.1 Hz) and that filtering data to this range discards potential differences in functional connectivity (Zuo et al., 2010; Kalcher et al., 2014; Gohel and Biswal, 2015; Wang et al., 2018). Accordingly, we followed a frequency-based approach, which yielded two main insights. First, there was a meaningful subdivision of COn in Slow-3 into its two core regions (e.g., Seeley et al., 2007), the insula and the ACC. This subdivision showed a particular inter-FC pattern, with significant inter-FC of the *insula*-subcomponent with visual networks and significant inter-FC of the *ACC*-subcomponent with RFPn. Of note, this subdivision only occurred in Slow-3 (also see Salvador et al., 2008), which includes frequencies typically filtered out (e.g., > 0.1), and is in agreement with task-fMRI evidence of a functional dissociation between the insula (alerting) and the ACC (set switching) (Han et al., 2019) and previous rs-fMRI evidence of the intrinsic connectivity of the (posterior dorsal) insula with visual brain areas (Cauda et al., 2011).

Second, the inter-FC of COn, RFPn, and visual networks appears stronger in some frequencies (Figure 4), though it is present across all frequency bins (Figure 3) (see also, Gohel and Biswal, 2015). For COn, the strongest inter-FC with visual networks was found in Slow-3 (faster than the typically analyzed rs-fMRI BOLD signal frequency range), whereas for RFPn, the strongest inter-FC with visual networks was found in Slow-5 and Slow-4 (the typically analyzed frequency range), aligning well with previous evidence on brain-regional differences in frequency power (e.g., Baria et al., 2011). The strongest inter-FC between visual network pairs was found in Slow-5, in accordance with previous documentations that the fMRI-BOLD signal in occipital networks mostly concentrates in infra-slow frequencies (i.e., 0.01-0.06 Hz; Wu et al., 2008; Baria et al., 2011) and correlates positively with lower-frequency electroencephalographic amplitude (i.e., delta and theta rhythms; ∼1.0–8.2 Hz; Jann et al., 2010). From a methods perspective, these results, along with the found association between the inter-FC of RFPn with visual networks in Slow-2 and VPS parameter *C*, provide supportive evidence that a frequency-binned approach can yield additional, valuable insights into the rs-fMRI BOLD signal.

In interpreting our results, some limitations should be considered. First, the two COn subcomponents were identified only in Slow-3. Future studies should determine whether this separation is specific to Slow-3. Further, the results on the strongest inter-FC in specific frequency bins do not directly imply ‘communication’ preferences between the neuronal populations of those networks. Rather, inter-FC was observed across the entire spectrum measured with BOLD-fMRI, which is in line with electrophysiological evidence showing that the oscillations that characterize functional networks span multiple frequency bands (Mantini et al., 2007). Finally, task-based fMRI studies investigating VPS variability *within* an individual could help elucidate the meaning and relevance of the observed negative association between inter-FC (of RFPn and visual networks) and VPS. Despite its limitations, our study underscores the usefulness of a frequency-based approach for better understanding the spontaneous fMRI-BOLD activity and how it links to behavior (Wu et al., 2008; Kalcher et al., 2014; Sasai et al., 2014).

## Conclusion

Our study provides first empirical evidence that the intrinsic inter-FC between RFPn and visual networks links to VPS. Albeit expected, we did not find such a link for COn. Our frequency-based approach revealed that the inter-FC between functional networks relevant for VPS and their link to VPS are also observed in frequencies above 0.1 Hz.

## Acknowledgments

This work was supported by grants of the German Forschungsgemeinschaft to K.F. [grant number FI 1424/2e1] and C.S. [grant number SO 1336/1e1]; the European Union’s Framework Programme for Research and Innovation Horizon 2020 (2014-2020) under the Marie Skłodowska-Curie Grant Agreements No. 859890 (SmartAge) and No. 754388 (LMUResearchFellows); and from LMUexcellent, funded by the Federal Ministry of Education and Research (BMBF) and the Free State of Bavaria under the Excellence Strategy of the German Federal Government and the Länder. We thank Dr. Mario E. Archila-Meléndez for helpful feedback on an earlier version of this paper.

## Abbreviations

ACC: anterior cingulate cortex
BOLD: Blood oxygenation level-dependent signal
COn: cingulo-opercular network
IC: independent component
ICA: independent component analysis
inter-FC: between-network functional connectivity
RFPn: right frontoparietal network
rs-fMRI: resting-state functional magnetic resonance imaging
TVA: theory of visual attention
VPS: visual processing speed

## References

Allen EA et al. (2011) A Baseline for the Multivariate Comparison of Resting-State Networks. Front Syst Neurosci 5

Ashburner J (2007) A fast diffeomorphic image registration algorithm. NeuroImage 38:95–113.

Baria AT, Baliki MN, Parrish T, Apkarian AV (2011) Anatomical and Functional Assemblies of Brain BOLD Oscillations. J Neurosci 31:7910–7919.

Beall EB, Lowe MJ (2007) Isolating physiologic noise sources with independently determined spatial measures. NeuroImage 37:1286–1300.

Beck AT, Steer RA, Brown G (1996) Beck depression inventory–II. Psychol Assess.

Beckmann CF, Mackay CE, Filippini N, Smith SM (2009) Group comparison of resting-state FMRI data using multi-subject ICA and dual regression. Neuroimage 47:S148.

Benjamini Y, Hochberg Y (1995) Controlling the false discovery rate: a practical and powerful approach to multiple testing. J R Stat Soc Ser B Methodol 57:289–300.

Bundesen C (1990) A theory of visual attention. Psychol Rev 97:523–547.

Bundesen C, Habekost T, Kyllingsbæk S (2005) A Neural Theory of Visual Attention: Bridging Cognition and Neurophysiology. Psychol Rev 112:291–328.

Cauda F, D’Agata F, Sacco K, Duca S, Geminiani G, Vercelli A (2011) Functional connectivity of the insula in the resting brain. NeuroImage 55:8–23.

Chechlacz M, Gillebert CR, Vangkilde SA, Petersen A, Humphreys GW (2015) Structural Variability within Frontoparietal Networks and Individual Differences in Attentional Functions: An Approach Using the Theory of Visual Attention. J Neurosci 35:10647–10658.

Cordes D, Haughton VM, Arfanakis K, Carew JD, Turski PA, Moritz CH, Quigley MA, Meyerand ME (2001) Frequencies Contributing to Functional Connectivity in the Cerebral Cortex in “Resting-state” Data. Am J Neuroradiol 22:1326–1333.

Crittenden BM, Mitchell DJ, Duncan J (2016) Task Encoding across the Multiple Demand Cortex Is Consistent with a Frontoparietal and Cingulo-Opercular Dual Networks Distinction. J Neurosci 36:6147–6155.

Dosenbach NUF, Fair DA, Cohen AL, Schlaggar BL, Petersen SE (2008) A dual-networks architecture of top-down control. Trends Cogn Sci 12:99–105.

Dosenbach NUF, Fair DA, Miezin FM, Cohen AL, Wenger KK, Dosenbach RAT, Fox MD, Snyder AZ, Vincent JL, Raichle ME, Schlaggar BL, Petersen SE (2007) Distinct brain networks for adaptive and stable task control in humans. Proc Natl Acad Sci 104:11073–11078.

Dyrholm M, Kyllingsbæk S, Espeseth T, Bundesen C (2011) Generalizing parametric models by introducing trial-by-trial parameter variability: The case of TVA. J Math Psychol 55:416–429.

Finke K, Bublak P, Krummenacher J, Kyllingsbæk S, Müller HJ, Schneider WX (2005) Usability of a theory of visual attention (TVA) for parameter-based measurement of attention I: Evidence from normal subjects. J Int Neuropsychol Soc 11

Gilbert CD, Li W (2013) Top-down influences on visual processing. Nat Rev Neurosci 14:350–363.

Gohel SR, Biswal BB (2015) Functional Integration Between Brain Regions at Rest Occurs in Multiple-Frequency Bands. Brain Connect 5:23–34.

Green JJ, McDonald JJ (2008) Electrical Neuroimaging Reveals Timing of Attentional Control Activity in Human Brain. PLOS Biol 6:e81.

Han SW, Eaton HP, Marois R (2019) Functional Fractionation of the Cingulo-opercular Network: Alerting Insula and Updating Cingulate. Cereb Cortex 29:2624–2638.

Jann K, Koenig T, Dierks T, Boesch C, Federspiel A (2010) Association of individual resting state EEG alpha frequency and cerebral blood flow. NeuroImage 51:365–372.

Jenkinson M, Beckmann CF, Behrens TEJ, Woolrich MW, Smith SM (2012) FSL. NeuroImage 62:782–790.

Kalcher K, Boubela RN, Huf W, Bartova L, Kronnerwetter C, Derntl B, Pezawas L, Filzmoser P, Nasel C, Moser E (2014) The Spectral Diversity of Resting-State Fluctuations in the Human Brain. PLOS ONE 9:e93375.

Kyllingsbæk S (2006) Modeling visual attention. Behav Res Methods 38:123–133.

Lehrl S, Merz J, Burkhard G, Fischer S (1999) Mehrfachwahl-Wortschatz-Intelligenztest. MWT-B 4th Ed Bal Spitta.

Mantini D, Perrucci MG, Gratta CD, Romani GL, Corbetta M (2007) Electrophysiological signatures of resting state networks in the human brain. Proc Natl Acad Sci 104:13170–13175.

Matsui T, Murakami T, Ohki K (2016) Transient neuronal coactivations embedded in globally propagating waves underlie resting-state functional connectivity. Proc Natl Acad Sci 113:6556–6561.

McAvinue LP, Habekost T, Johnson KA, Kyllingsbæk S, Vangkilde S, Bundesen C, Robertson IH (2012) Sustained attention, attentional selectivity, and attentional capacity across the lifespan. Atten Percept Psychophys 74:1570–1582.

Penttonen M, Buzsáki G (2003) Natural logarithmic relationship between brain oscillators. Thalamus Relat Syst 2:145–152.

Power JD, Barnes KA, Snyder AZ, Schlaggar BL, Petersen SE (2012) Spurious but systematic correlations in functional connectivity MRI networks arise from subject motion. NeuroImage 59:2142–2154.

Preibisch C, G JGC, Bührer M, Riedl V (2015) Evaluation of Multiband EPI Acquisitions for Resting State fMRI. PLOS ONE 10:e0136961.

R Core Team (2020) R: The R Project for Statistical Computing. Available at: https://www.r-project.org/.

Raichle ME (2011) The Restless Brain. Brain Connect 1:3–12.

Reitan RM (1958) Validity of the Trail Making Test as an Indicator of Organic Brain Damage. :6.

Riedl V, Utz L, Castrillón G, Grimmer T, Rauschecker JP, Ploner M, Friston KJ, Drzezga A, Sorg C (2016) Metabolic connectivity mapping reveals effective connectivity in the resting human brain. Proc Natl Acad Sci 113:428–433.

Ries A, Chang C, Glim S, Meng C, Sorg C, Wohlschläger A (2018) Grading of Frequency Spectral Centroid Across Resting-State Networks. Front Hum Neurosci 12.

Ruiz-Rizzo AL, Küchenhoff S (2020) Inter-FC. OSF. Available at: https://osf.io/nhqg3/.

Ruiz-Rizzo AL, Neitzel J, Müller HJ, Sorg C, Finke K (2018) Distinctive Correspondence Between Separable Visual Attention Functions and Intrinsic Brain Networks. Front Hum Neurosci 12.

Ruiz-Rizzo AL, Sorg C, Napiórkowski N, Neitzel J, Menegaux A, Müller HJ, Vangkilde S, Finke K (2019) Decreased cingulo-opercular network functional connectivity mediates the impact of aging on visual processing speed. Neurobiol Aging 73:50–60.

Sadaghiani S, Scheeringa R, Lehongre K, Morillon B, Giraud A-L, D’Esposito M, Kleinschmidt A (2012) Alpha-Band Phase Synchrony Is Related to Activity in the Fronto-Parietal Adaptive Control Network. J Neurosci 32:14305–14310.

Salvador R, Martínez A, Pomarol-Clotet E, Gomar J, Vila F, Sarró S, Capdevila A, Bullmore E (2008) A simple view of the brain through a frequency-specific functional connectivity measure. NeuroImage 39:279–289.

Sasai S, Homae F, Watanabe H, Sasaki AT, Tanabe HC, Sadato N, Taga G (2014) Frequency-specific network topologies in the resting human brain. Front Hum Neurosci 8.

Scolari M, Seidl-Rathkopf KN, Kastner S (2015) Functions of the human frontoparietal attention network: Evidence from neuroimaging. Curr Opin Behav Sci 1:32–39.

Seeley WW, Menon V, Schatzberg AF, Keller J, Glover GH, Kenna H, Reiss AL, Greicius MD (2007) Dissociable Intrinsic Connectivity Networks for Salience Processing and Executive Control. J Neurosci 27:2349–2356.

Sestieri C, Corbetta M, Spadone S, Romani GL, Shulman GL (2013) Domain-general Signals in the Cingulo-opercular Network for Visuospatial Attention and Episodic Memory. J Cogn Neurosci 26:551–568.

Smith SM, Jenkinson M, Woolrich MW, Beckmann CF, Behrens TEJ, Johansen-Berg H, Bannister PR, De Luca M, Drobnjak I, Flitney DE, Niazy RK, Saunders J, Vickers J, Zhang Y, De Stefano N, Brady JM, Matthews PM (2004) Advances in functional and structural MR image analysis and implementation as FSL. NeuroImage 23:S208–S219.

Sperling G (1960) The information available in brief visual presentations. Psychol Monogr Gen Appl 74:1–29.

Sridharan D, Levitin DJ, Menon V (2008) A critical role for the right fronto-insular cortex in switching between central-executive and default-mode networks. Proc Natl Acad Sci 105:12569–12574.

Thiebaut de Schotten M, Dell’Acqua F, Forkel S, Simmons A, Vergani F, Murphy DGM, Catani M (2011) A Lateralized Brain Network for Visuo-Spatial Attention. Nat Preced:1–1.

Tombaugh TN (2004) Trail Making Test A and B: Normative data stratified by age and education. Arch Clin Neuropsychol 19:203–214.

Uddin LQ, Yeo BTT, Spreng RN (2019) Towards a Universal Taxonomy of Macro-scale Functional Human Brain Networks. Brain Topogr 32:926–942.

Wang Y, Zhu L, Zou Q, Cui Q, Liao W, Duan X, Biswal B, Chen H (2018) Frequency dependent hub role of the dorsal and ventral right anterior insula. NeuroImage 165:112–117.

Wu CW, Gu H, Lu H, Stein EA, Chen J-H, Yang Y (2008) Frequency specificity of functional connectivity in brain networks. NeuroImage 42:1047–1055.

Yan C, Zang Y (2010) DPARSF: a MATLAB toolbox for “pipeline” data analysis of resting-state fMRI. Front Syst Neurosci 4.

Zuo X-N, Di Martino A, Kelly C, Shehzad ZE, Gee DG, Klein DF, Castellanos FX, Biswal BB, Milham MP (2010) The oscillating brain: Complex and reliable. NeuroImage 49:1432–1445.

